# Chromatin reorganization during myoblast differentiation involves the caspase-dependent removal of SATB2

**DOI:** 10.1101/2019.12.19.883579

**Authors:** Ryan A.V. Bell, Mohammad H. Al-Khalaf, Steve Brunette, Alphonse Chu, Georg Dechant, Galina Apostolova, Jeffrey Dilworth, Lynn A. Megeney

## Abstract

Induction of lineage-specific gene programs are strongly influenced by alterations in local chromatin architecture. However, key players that impact this genome reorganization remain largely unknown. Here, we report that removal of special AT-rich binding protein 2 (SATB2), a nuclear protein that binds matrix attachment regions, is a key event in initiating myogenic differentiation. Deletion of SATB2 in muscle cell culture models and in vivo, accelerates differentiation and depletes the muscle progenitor pool, respectively. Genome wide analysis indicates that SATB2 binding is both repressive and inductive, as loss of SATB2 leads to expression of differentiation regulatory factors and inhibition of genes that impair this process. Finally, we noted that the differentiation-specific decline in SATB2 protein is dependent on a caspase 7-mediated cleavage event. Taken together, this study demonstrates that temporal control of SATB2 protein is critical for shaping the chromatin environment and coordinating the myogenic differentiation program.

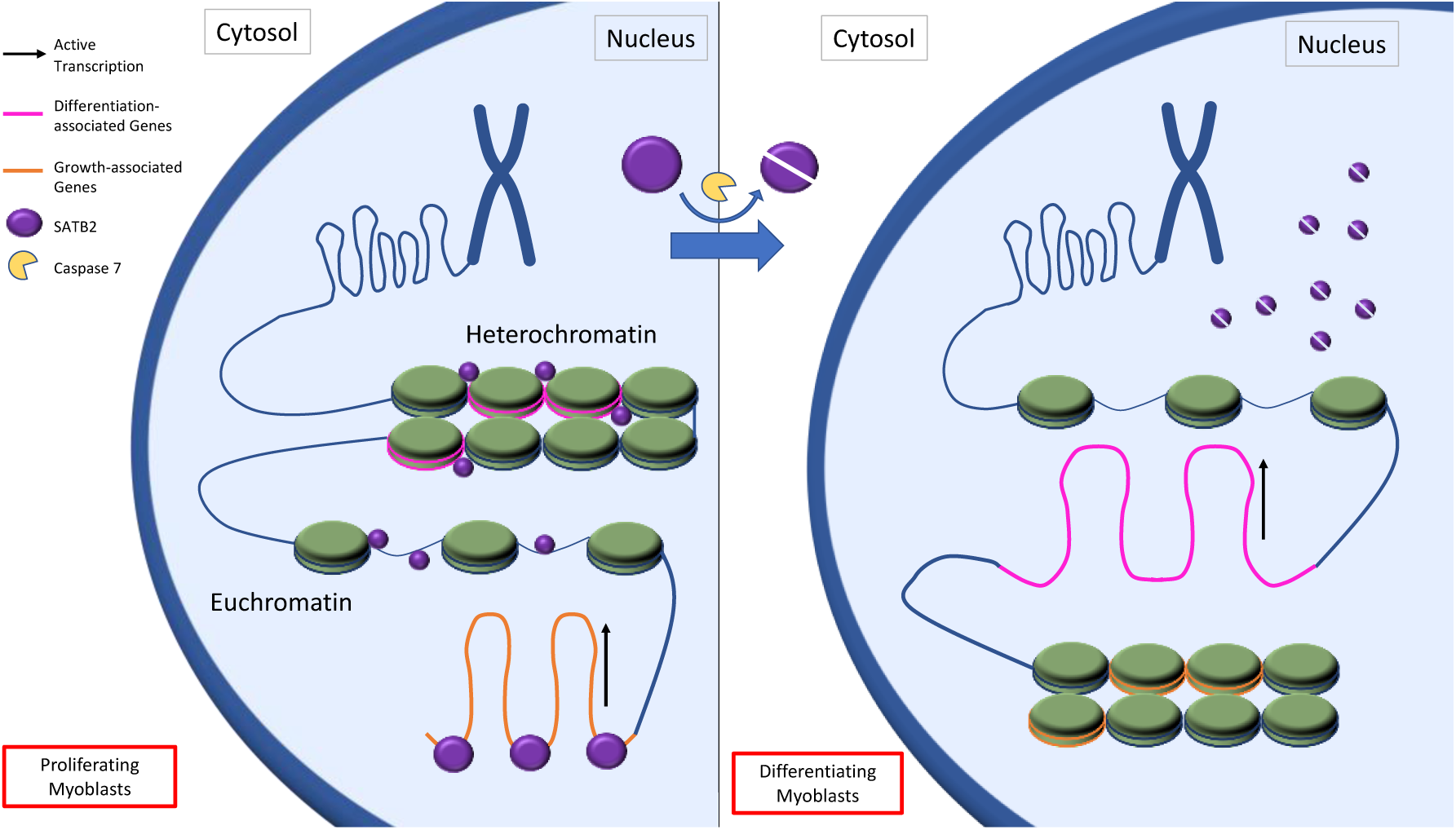
Graphical Abstract

## Introduction

One of the key alterations that characterize a cell’s progression through differentiation is a restructuring of the nuclear landscape to allow for the expression of lineage-specific genes. Indeed, chromatin conformation undergoes significant restructuring when cells progress from replicating to differentiating phenotypes, with changes occurring to repress or open access at specific gene loci (Müller and Leutz, 2001; Forcales et al. 2012). These genome alterations are facilitated by two general mechanisms, loci-specific modifications of DNA and histones and targeting of structural proteins that control the higher-order structure of chromatin (Gómez-Díaz and Corces, 2014; Hota and Bruneau, 2016). Progress through the myogenic differentiation program is no exception to this paradigm, with multiple genomic conversions from heterochromatin to euchromatin during myogenesis to allow for muscle-specific genes to be expressed (Doynova et al. 2017). A number of proteins that govern DNA and histone methylation changes have been implicated in the control of myogenesis (reviewed by Robinson and Dilworth, 2018), yet little information exists on the role of proteins that manage higher-order chromatin reorganization during this cell fate determining phase.

The special AT-rich binding proteins (SATB1 and SATB2) are a family of matrix attachment region (MAR) binding proteins that are commonly expressed in stem cells (Savarese et al., 2009; Dong et al., 2015) and play an active role in the global organization of chromatin. MAR binding proteins are unique in that they mediate both repressive and inductive signals for gene expression. This duality of MAR protein function depends on the proximity of gene promoters and insulator regions relative to the position where the MAR protein anchors the DNA to the chromatin scaffold (Bushey et al., 2008; Arope et al., 2013). There is increasing evidence that these structural proteins, particularly SATB2, are vital to the epigenetic regulation of numerous genes, several of which are involved in stem cell fate determination and maintaining cancer cell progression (Britanova et al., 2005; Dobreva et al., 2006; Han et al., 2008; Savarese et al., 2009; Dong et al., 2015; Leone et al., 2015). Moreover, human *SATB2* mutations are associated with SATB2 syndrome, which is characterized by a plethora of developmental abnormalities, including facial dysmorphia, neural and bone defects, and weak muscle tone in infancy (Bengani et al. 2017). Given these observations, regulation of SATB2 protein content may be a critical step in managing chromatin ultrastructure and gene expression during cell maturation in general.

There is no information regarding how SATB2 manages such a diverse biologic response, nor how SATB2 protein is directed to or removed from its associated genomic targets. However, caspase-mediated cleavage of SATB1 has been shown to be an important step in the control of gene expression that proceeds T-cell apoptosis, where cleavage of SATB1 results in its release from MARs (Galande et al. 2001). Interestingly, caspase proteases have prominent roles in non-death processes such as differentiation, inflammation, remodeling, and cell survival (Unsain and Parker, 2015). The role of caspases in differentiation is particularly well established, with both initiator and executioner caspases being involved in the development of a variety of tissues (reviewed in Bell and Megeney, 2017). With respect to skeletal muscle differentiation, catalytically active caspase 3 and 9 appear to be required for progenitor progression through the myogenic program (Fernando et al. 2002; Murray et al., 2008; Larsen et al. 2010).

The integral nature of caspase activity during myoblast differentiation and the possibility that caspases could act on chromatin organizing proteins, led us to investigate the behavior of SATB2 within muscle progenitor cells and the potential that caspase enzymes mediate its function.

## Results

### SATB2 restrains induction of muscle cell differentiation

Western blot and immunofluorescent analyses identified SATB2 as a nuclear protein within proliferating C2C12 muscle cells, which was markedly reduced in expression during the early stages of differentiation (Figure 1A and B). When expressed, SATB2 was mainly relegated to the euchromatic nuclear space, however there was a subfraction of SATB2 co-staining with heterochromatin protein 1 alpha (HP1α) occurring in the more condensed regions of the nucleus (Figure 1C). Having established that SATB2 protein is reduced during myogenesis, we next sought to determine the role of SATB2 in muscle cell proliferation and differentiation. To this end, we initially designed CRISPR-Cas9 guide RNAs to inhibit SATB2 gene expression in replicating myoblasts. However, infection of C2C12 muscle cells with an adenovirus expressing enhanced Cas9 (Slaymaker et al., 2016) led to a dramatic increase in SATB2 protein content in post-differentiated myoblasts, at a time when SATB2 would otherwise be in significant decline (see Supplementary Figure 1). Given this unexpected impact of Cas9 on our protein of interest we chose to pursue short interfering RNA (siRNA)-mediated targeting of SATB2 as a means to address its biologic role in cell culture models. siRNA targeting of SATB2 (siSATB2) did not produce any noticeable alteration in cell viability or growth (Supplementary Figure 2). However, siSATB2-treated cells displayed enhanced differentiation kinetics (Figure 1D) resulting in a marked increase in myosin heavy chain (MHC) expression at the time points tested compared to cells treated with scrambled siRNA (siControl) (Figure 1E). Moreover, the formation of multinucleated myotubes was dramatically accelerated in SATB2 targeted cells compared to the control condition (Figure 1F). These results indicate that SATB2 plays an important role in mediating the transition from proliferation to induction of the myogenic differentiation program.

**Figure 1.**
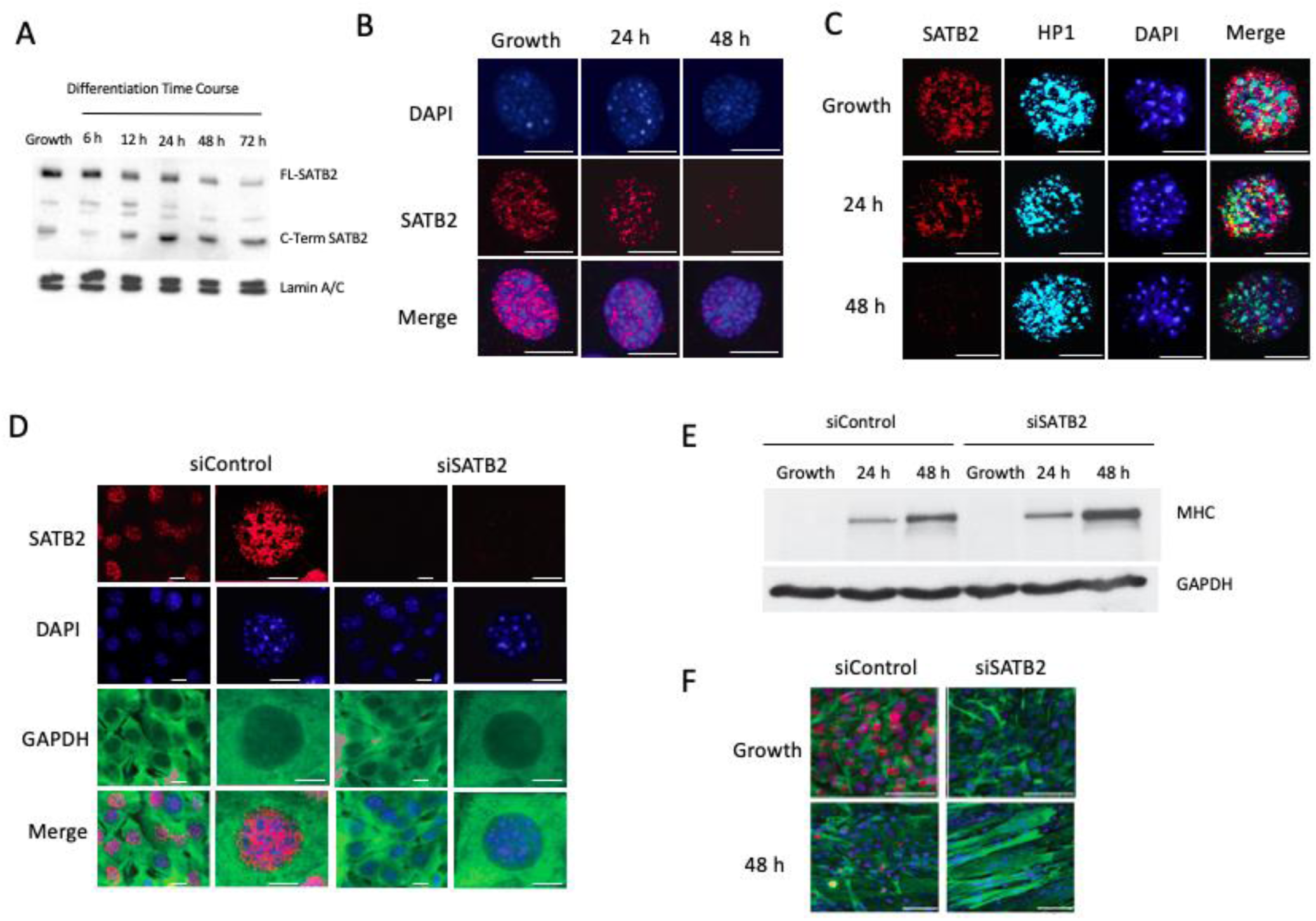
SATB2 regulates myogenic gene expression during myoblast differentiation. (A) Representative western blot showing the decreased expression of full-length SATB2 (FL-SATB2) during a differentiation time course for C2C12 cells. This was accompanied by a concomitant increase in a C-terminal fragment of SATB2 (C-Term SATB2). (B) Representative immunofluorescent staining of SATB2 in proliferating and differentiated C2C12 cells. Images are representative of n = 5 independent samples at each time point. DAPI (blue), SATB2 (red); scale = 10 μM. (C) Representative immunofluorescent images depicting SATB2 localization in relation to heterochromatin protein 1α (HP1α). Images are representative from n = 3 independent determinations for each time point. Scale bar = 10 μM. DAPI (blue), SATB2 (red), HP1α (teal). (D) Immunofluorescent images depicting the efficacy of SATB2 depletion using siRNA. Muscle cell cytoplasm was counterstained with an anti-GAPDH antibody. Images are representative of n = 3 independent determinations. DAPI (blue), GAPDH (green), and SATB2 (red). Scale bar for the low magnification column = 10 μM. Scale bar for the higher magnification column = 5 μM. (E) Western blot showing the myosin heavy chain (MHC) protein expression during growth and differentiation time points following siControl or siSATB2 treatments in C2C12 cells. Image is representative of n = 3 independent determinations. GAPDH was used as a loading control. (F) Immunofluorescent images indicating that the depletion of SATB2 leads to early myotube formation after C2C12 cells were induced to differentiate. Images are representative of n = 3 independent determinations. DAPI (blue), SATB2 (red), and desmin (green). Scale bar = 50 μM.

### In vivo depletion of SATB2 in muscle satellite cells decreases muscle fiber area and the number of Pax7-expressing satellite cells

To examine whether loss of SATB2 would alter the muscle cell differentiation program in vivo, we generated satellite cell-specific deletion of SATB2, under the control of tamoxifen induction, by crossing the *Satb2_fl/fl_* mouse strain (Jaitner et al., 2016) with the *Pax7CreER* mouse strain (Nishijo et al., 2009) to generate *Pax7CreER/Satb2_fl/fl_* mice. *Pax7CreER/Satb2_fl/fl_* mice (and requisite controls, *Satb2_fl/fl_* strain) were given tamoxifen at three weeks of age and monitored continuously (as per University of Ottawa Animal Care and Veterinary Service (ACVS) guidelines). Gross assessment of motor function and physical state indicated that there were no substantial motor effects stemming from the depletion of SATB2. Quantitative assessment of mice weights showed no significant differences between control and *Pax7CreER/Satb2_fl/fl_* mice (data not shown). However, histologic/morphologic analysis revealed notable alterations in skeletal muscle structure between control and *Pax7CreER/Satb2_fl/fl_* mice. For example, measurement of fiber area within the tibialis anterior (TA) from control and *Pax7CreER/Satb2_fl/fl_* mice indicated that the removal of SATB2 from muscle satellite cells led to a significant decrease in fiber area concurrent to an increase in fiber number as compared to the control strains (Figure 2A–C). Moreover, following tamoxifen treatment, the number of Pax7-expressing satellite cells within *Pax7CreER/Satb2_fl/fl_* TA muscle decreased substantially when compared to the *Satb2_fl/fl_* control muscle (Figure 2D). Collectively, these observations are consistent with the hypothesis that loss of SATB2 in the satellite cell niche leads to premature activation and differentiation of these muscle progenitor cells, which is reflected at the anatomical level by increased numbers of smaller muscle fibers.

**Figure 2.**
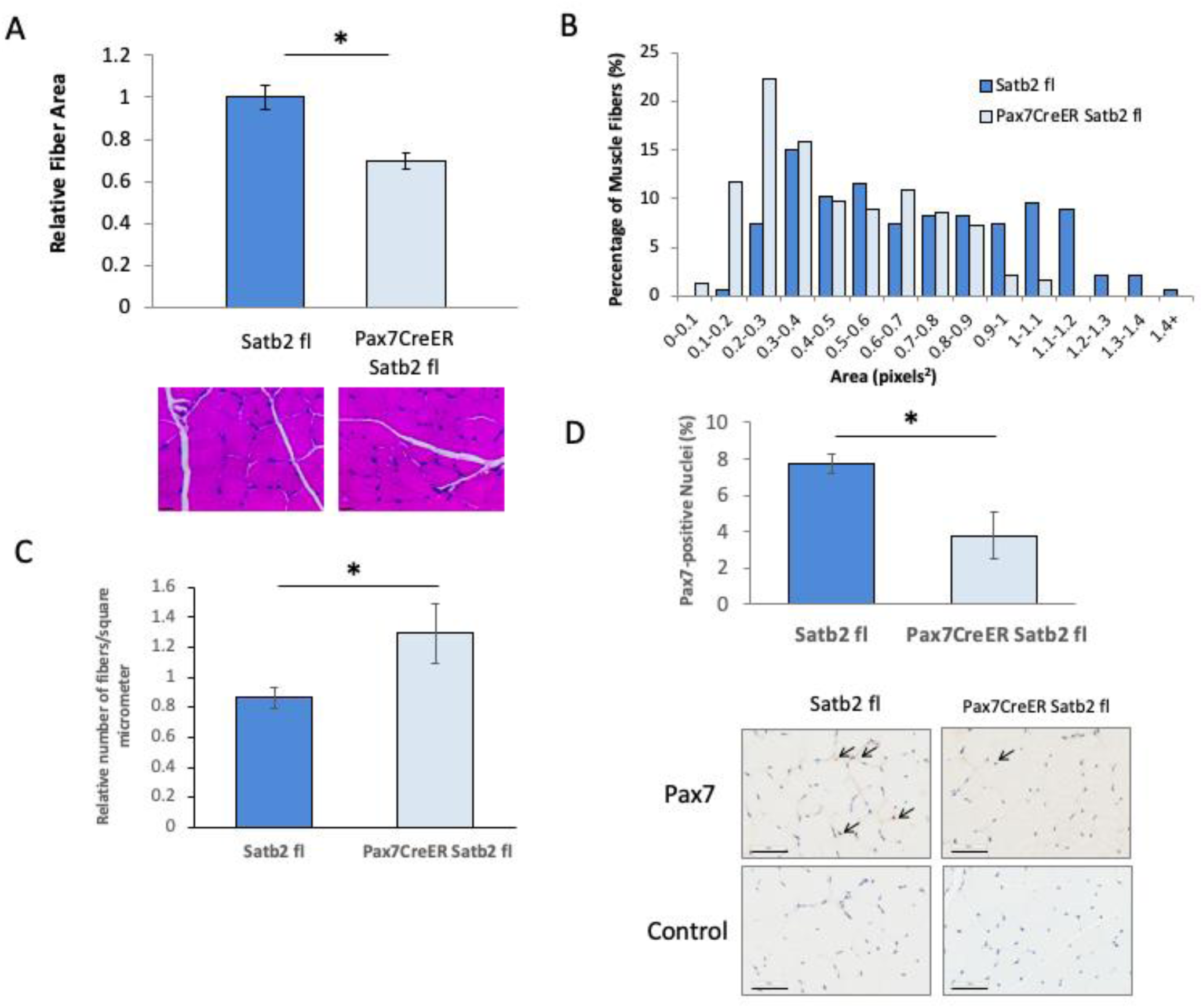
In vivo reduction of SATB2 in muscle satellite cells decreases muscle fiber area and the number of Pax7-expressing satellite cells. (A) Tibialis anterior fiber areas decreased in *Pax7CreER/Satb2_fl/fl_* mice as compared to control (*Satb2_fl/fl_*) mice. ImageJ was used to the measure the fiber areas of hematoxylin and eosin (H&E) stained tissue sections. Data are the means ± SEM, n = 3 independent determinations on separate tissue samples. H&E stained sections are representative of n = 3 independent determinations. Scale bar: 100 μM. (B) The distribution of muscle fiber sizes across 15 bins. Over 200 muscle fibers were measured for both control and *Pax7CreER/Satb2_fl/fl_* mice. (C) Bar graph indicating the significant increase in the relative number of muscle fibers per μM_2_. Data are the means ± SEM, n = 3 independent determinations on separate tissue samples. (D) The percentage of nuclei that were Pax7-positive in control (Satb2 fl) and Satb2-ablated (Pax7CreER Satb2 fl) mouse muscle. The panels below the bar graph are representative tissue sections stained with a Pax7 antibody (Pax7) or no antibody (Control); scale: 50 μM. Arrows indicate positively stained nuclei. Data are representative of n = 3 independent determinations on separate tissue samples. * indicates a significant difference between *Pax7CreER/Satb2_fl/fl_* and *Satb2_fl/fl_* mice as determined by the Student’s *t*-test (p < 0.05).

### Chromatin immunoprecipitation (ChIP)- and RNA-seq data support a role for SATB2 in regulating muscle satellite cell differentiation

In order to more clearly identify the genes bound and regulated by SATB2 during muscle cell differentiation, we performed both ChIP- and RNA-seq analyses. Our ChIP-seq data was subjected to Gene Ontology categorization using the GREAT software (McLean et al., 2010), which indicated that of the SATB2 binding peaks identified, 91.5% of the reads mapped to the genome (Figure 3A). This analysis revealed that SATB2 bound to a large proportion of the mouse genome, covering 38% of known classified genes (Figure 3B). Unsurprisingly, given that SATB2 is recognized as a structural DNA binding protein, SATB2 was found to bind to many genetic loci in proliferating C2C12 cells. The binding targets spanned a variety of biological processes (Figure 3C and Supplementary Table 1), with SATB2 bound most frequently to genes associated with chromatin or chromosome organization and cell division. This genome targeting behavior of SATB2 is consistent with the observations in Figures 1 and 2, suggesting that SATB2 may control gene expression during muscle cell proliferation and/or the transition of myoblasts from growth to differentiation.

**Figure 3.**
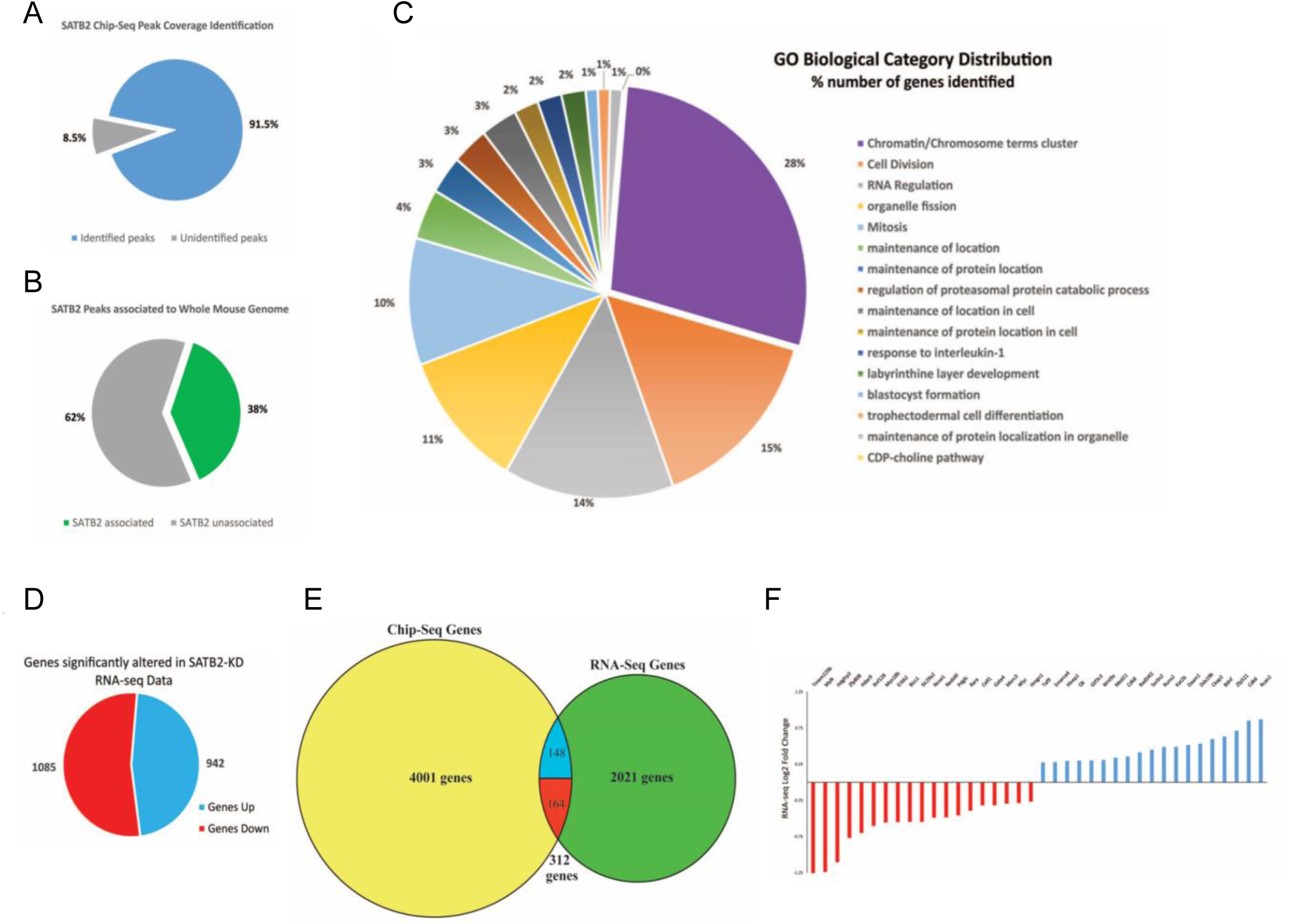
SATB2 regulates the expression of many genes associated with various biological processes within myoblasts. (A) GREAT-input identified 8413 (91.5%) of the 9010 peaks generated from SATB2 ChIP-sequencing; (B) identified peaks corresponded to 38% of the 21 902 *Mus musculus* genes classified in the UCSC mm10 NCBI build 38 species assembly. (C) Pie chart shows the distribution of the genes identified as it relates to Gene Ontology (GO) category clusters shown in Supplementary Table S1. (D) Cuffdiff analysis of the RNA-seq dataset show that 2027 genes were identified to have significantly altered expression between the control and knockdown conditions, with 1085 genes being significantly downregulated in expression and 942 genes being significantly upregulated. (E) Venn diagram comparing MACS-identified SATB2 ChIP-seq genes (4001) from C2C12 cells under growth conditions with Cuffdiff-identified SATB2 knockdown RNA-seq genes (2021) from 24 h-differentiated cells. Three hundred and twelve genes were found to be bound to SATB2 in proliferating cells, which was altered significantly following the reduction in *Satb2* gene expression. Of those 312 genes, 164 were significantly downregulated and 148 were significantly upregulated. (F) Graph highlights a selection from the 312 genes identified in common between the ChIP-seq and RNAseq datasets. These selected genes (19 upregulated and 19 downregulated) are all known to be involved in the genetic switch from a proliferative program to a differentiation program. The RNA-seq dataset is the result of the analysis of n = 3 independent determinations.

To characterize the impact of SATB2 on gene expression during early myoblast differentiation, we performed RNA-seq on siControl- and siSATB2-treated cells that had been differentiated for 24 h. Prior to the analysis of our RNA-seq dataset, we confirmed that siSATB2 was able to significantly reduce *Satb2* expression (Supplementary Figure 3). Assessment of the RNA expression profiles using CuffDiff2 (Trapnell et al. 2013) indicated that 2021 genes were significantly altered in the *Satb2*-depleted samples as compared to the control samples (Figure 3D and E). Of these genes, 942 were upregulated and 1085 were downregulated following the decrease in *Satb2* expression (Figure 3D). Three hundred and twelve of these genes had been previously identified in our ChIP-seq data set (Figure 3E), indicating a particular set of genes that was directly regulated by SATB2 binding. Of those 312 genes, approximately half (148) were upregulated while the other half was downregulated (Figure 3E). Figure 3F shows a subset of the genes identified by RNA-seq and ChIP-seq that indicate the potential role of SATB2 in regulating cell proliferation and differentiation pathways. For example, several of the targets with reduced expression include genes known to repress differentiation, such as *Hdac9, Ncoa1, Hdgfrp3, Rnf128,* and *Celf1*, whereas a large number of targets with enhanced expression are proteins known to accelerate differentiation, including *Smarca4*, *Wnt9a*, *Kat2b*, *Med21*, *Cdk8*, and *Bdnf*. This heterogenous binding and gene regulation activity of SATB2 is consistent with the euchromatin and heterochromatin distribution during growth and early stages of muscle cell differentiation, suggesting that SATB2 can promote and repress gene expression depending on the loci it targets.

### Caspase 7 cleaves SATB2 during early myogenesis

The mechanisms that govern chromatin remodeling during cellular differentiation remain largely unknown, but evidence from apoptotic nuclei suggests that nuclear structural proteins are targeted by caspase proteases (Galande et al., 2001; Sun et al., 2006; Dudek et al., 2018). Moreover, our identification that a major C-terminal fragment of SATB2 becomes more prominent during muscle cell differentiation suggests that a proteolytic event may target the SATB2 protein (Figure 1A and Supplementary Figure 4). Prior studies from our laboratory have shown that caspase 3 plays a prominent role in myogenesis, targeting and cleaving a number of substrate proteins to engage the differentiation program (Fernando et al. 2002; Larsen et al. 2010; Dick et al. 2015). To examine whether SATB2 proteolysis occurred, we suppressed endogenous effector caspase activity (caspase 3 and 7) with the peptide inhibitor z.DEVD.fmk and reassessed SATB2 protein expression during muscle cell differentiation. Caspase inhibition during differentiation led to sustained expression of SATB2 protein in the nucleus compared to control cells (Figure 4A and B). In an attempt to attribute SATB2 cleavage to either caspase 3 or 7 (or both), an in vitro cleavage assay was performed. This assay demonstrated that caspase 7 activity was very robust at targeting SATB2 protein, whereas caspase 3 did not induce measurable cleavage (Figure 4C). Mass spectrometry analysis of the caspase 7 mediated SATB2 fragments mapped a putative caspase cleavage site at D477 (Supplementary Figures 4 and 5).

**Figure 4.**
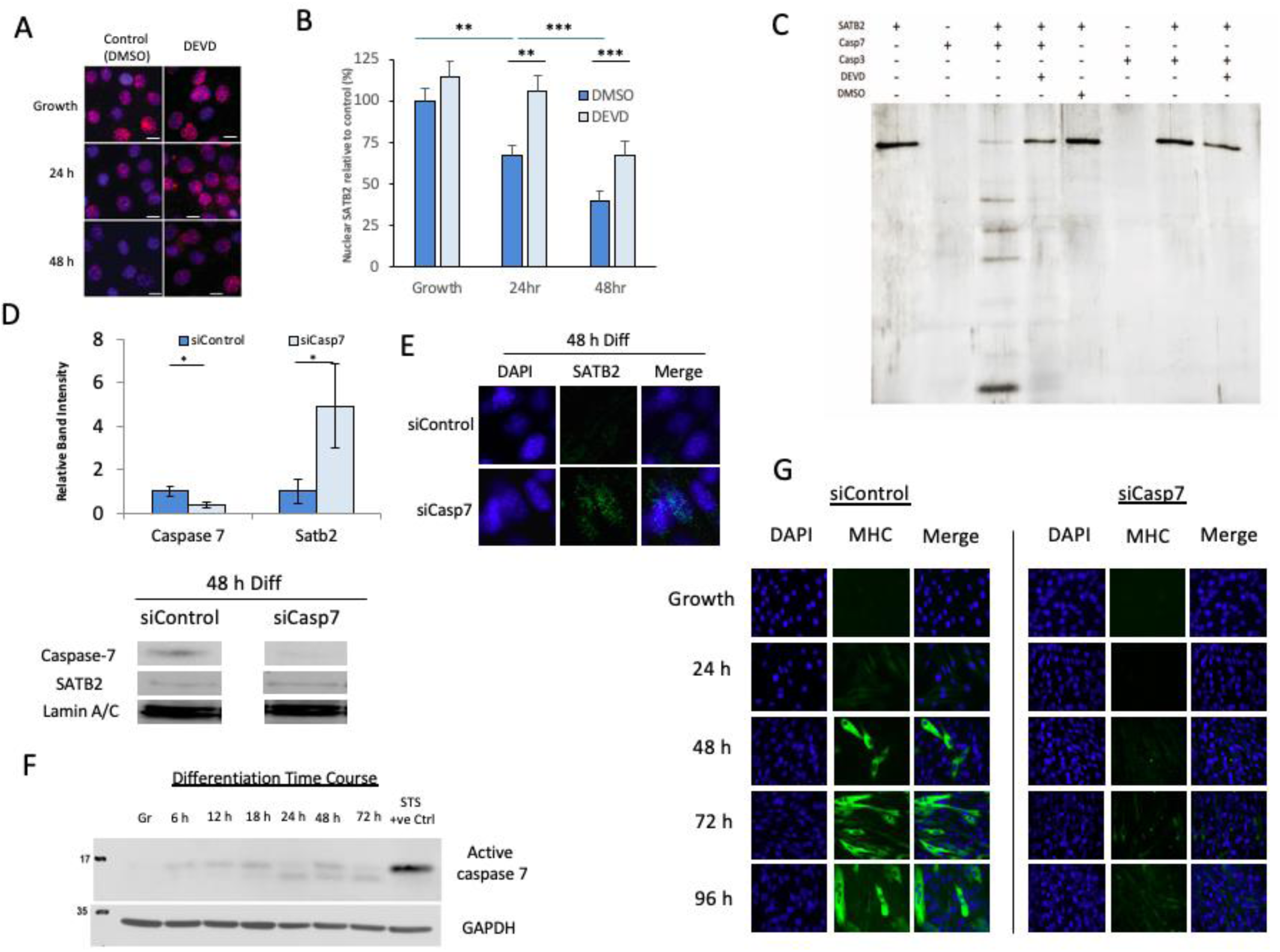
Caspase 7 cleaves SATB2 during early skeletal muscle differentiation and is necessary for myogenesis. (A) Immunofluorescent staining of SATB2 following caspase inhibition by z-DEVD.fmk (DEVD) in proliferating and differentiating C2C12 cells. Images are representative of n = 3 determinations on independent samples. DAPI (blue); SATB2 (red); scale: 10 μM. (B) Quantification of SATB2 nuclear expression following DEVD-mediated suppression of caspase activity. Data are the means ± SEM from n = 3 independent samples; 75–100 cells per condition/time point were analyzed. ** indicates p < 0.01 and *** indicates p < 0.001 as determined by the Student’s *t*-test. (C) Silver stained gel of an in vitro caspase cleavage assay. Reactions included combinations of recombinant SATB2 protein, active caspase 3/7 recombinant proteins, caspase chemical inhibitor (DEVD), or the control chemical DMSO. (D) Caspase 7 and SATB2 expression following siCasp7 and siControl treatments of C2C12 cells. Both the histogram and the representative western blots (n = 3) were the expression levels of those two proteins 48 h post-induction of differentiation. * indicates a significant difference in caspase 7 or SATB2 expression as determined by the Student’s *t*-test (p < 0.05) (E) Immunofluorescent images showing the sustained expression of SATB2 after the depletion of caspase 7 in C2C12 cells differentiated for 48 h. Images are representative of n = 3 independent determinations. (F) Representative western blot depicting the cleaved active fragment of caspase 7 found in the nuclear fraction of C2C12 cells during their proliferative phase (Gr) and at various stages of differentiation (6–72 h post-induction; n = 3). Staurosporine (STS) was used as a positive control. (G) Representative western blot showing the reduction in caspase expression in differentiating C2C12 myoblasts (48 h post-induction; n = 3). See the Supplemental Information for the quantification of the knockdown of caspase 7 (Supplemental Figure S6). (E) Immunofluorescent staining of myosin heavy chain (MHC) in siControl- and siCasp7-treated C2C12 myoblasts under growth and differentiation conditions. Panel is representative of n = 3 determinations on independent samples. DAPI (blue), MHC (green).

To corroborate caspase 7 as a targeting protease of SATB2 and by inference as a regulatory enzyme that may promote muscle cell differentiation, we utilized siRNA to suppress caspase 7 expression. siRNA repression of caspase 7 (siCasp7) led to a concomitant accumulation of SATB2 protein during C2C12 muscle cell differentiation when compared to siControl-treated cells (Figure 4D and E). Investigating caspase 7 more closely, we determined that the expression of nuclear, active caspase 7 was inversely correlated with SATB2 expression, with nuclear caspase 7 levels increasing during early myoblast differentiation (Figure 4F). To assess the broad effect of this protease on muscle differentiation, we compared siCasp7 cells (where siRNA treatment reduced caspase 7 expression by ∼60%; Supplementary Figure 6) vs. siControl-treated cells and noted that siCasp7 cultures displayed a significant impairment in low serum induction of differentiation, with a dramatic reduction in the expression of MHC and a near complete inhibition of multi-nucleate myotube formation (Figure 4G). Taken together, these results suggest that a signaling pathway engages caspase 7 activation to target SATB2 protein and that this intersection between a protease and MAR proteins is a critical determinant in securing the muscle cell differentiation program.

## Discussion

Chromatin remodeling plays a central role in stem cell differentiation, as it facilitates the dramatic shift in gene expression profiles that accompanies the exit from the cell cycle and the commitment to a particular lineage (Müller and Leutz, 2001; de la Serna et al. 2006, Keenen and de la Serna, 2008; Fisher and Fisher, 2011; Chen and Dent 2014; Dixon et al. 2015; Hota and Bruneau 2016; Ye et al., 2017). These changes are dependent upon both transcription mediated change concurrent with a shift in the epigenetic landscape, which may allow or limit gene expression in the relevant regions of the genome. One mechanism that will influence lineage-dependent transcription and epigenetic change is the presence (or absence) of higher order MAR proteins that mediate chromatin structure, and thereby physical access to key genetic loci (Hawkins et al., 2001; Savarese et al., 2009; Asanoma et al., 2012). While changes to the chromatin architecture within skeletal muscle progenitor cells have been observed (Doynova et al. 2017), the exact players that govern these changes are not well known. Here, we identified SATB2 as a chromatin organizer that plays a key role in mediating the progression of myoblasts into the myogenic differentiation program.

Interestingly, reduction of SATB2 expression did not affect myoblast proliferation (Supplementary Figure 2), indicating that SATB2 may primarily block cells from prematurely entering into a terminally differentiated state by denying access to important myogenic genes. Reduced expression of SATB2 during early myogenesis appears to accelerate the differentiated phenotype, as evidenced by the siRNA-mediated knockdown of SATB2, which hastens the expression of MHC (Figure 1E) and accelerates the formation of myotubes during early differentiation (Figure 1F). This coincides with our supposition that SATB2 may sequester myogenic genes (perhaps within the heterochromatic areas of the nucleus), and once removed, primes myoblasts for myogenic gene expression.

As noted above, SATB2 protein within muscle cells is mainly relegated to euchromatic regions of the nucleus, with a reduced but defined sub-localization of the protein in heterochromatic areas (Figure 1C). This localization pattern is evident during growth conditions and is consistent with the concept that SATB2 promotes cell proliferation through its action on genetic loci in euchromatin regions. The corollary to this is the impact on SATB2 targeted loci during differentiation, where SATB2 is removed from the nucleus, which simultaneously represses growth-related euchromatic genes and relieves repression of differentiation-specific genes that were previously sequestered in the heterochomatic regions of the nucleus. This hypothesis is supported by the ChIP- and RNA-seq data where a number of genes that are potent promoters of cell proliferation were downregulated during early myoblast differentiation, including *Hdac9* (Lapierre et al. 2016), *Celf1* (Xia et al., 2015), *Ncoa1* (Wang et al., 2018), *Hdgfrp3* (Xiao et al., 2013), and *Rnf128* (Lee et al., 2016), all of which were significantly downregulated following SATB2 removal during early stages of the differentiation program (Figure 3F). Conversely, we identified several differentiation-specific genes that showed an increase in expression following SATB2 depletion during early myogenesis. This list includes *Smarca4*, which is known to play a role in neurogenesis (Yu et al., 2013), erythropoiesis (Griffin et al., 2008), and myogenesis (Albini et al., 2015); *Wnt9a*, which is a positive regulatory factor in hematopoiesis (Richter et al., 2018) and mediates the transition from proliferation to differentiation in skeletal myoblasts (Tanaka et al., 2011). Furthermore, *Cdk8* was upregulated following the loss of SATB2 protein during early muscle differentiation, and is known to be important for mediating Notch degradation, which is critical for satellite cells to exit the cell cycle and enter the myogenic differentiation program (Buas and Kadesch, 2010). These findings support the premise that SATB2 in proliferating myoblasts acts to maintain cell proliferation, while the induction of differentiation leads to the loss of SATB2 and the repression of growth-associated genes and the activation of previously sequestered differentiation-associated gene program.

Caspase signaling is an inductive cue for skeletal muscle differentiation and a number of caspase 3 substrates have been identified which participate in the differentiation process. These proteins include the mammalian sterile twenty-like kinase, MST1, which when cleaved by caspase 3 engages a self-contained promyogenic signal (Fernando et al., 2002). Caspase 3 can also moderate earlier steps in the myogenic cascade by targeting proteins the promote stem cell self renewal such as Pax7 (Dick et al. 2015). Caspase 3 also engages a regulated form of DNA damage and repair, which acts a prodifferentiaon signal (Larsen et al. 2010; Al-Khalaf et al 2016). Here, caspase 3 engages activation of CAD by cleaving and removing its cognate inhibitor protein ICAD. Once activated CAD targets regions of the genome, inflicting strand breaks which activate expression of p21, a conserved cell cycle inhibitor that governs differentiation across multiple cell lineages (Larsen et al. 2010).

Despite the prominent role of caspase 3 in the differentiation process, we have noted that SATB2 is cleaved exclusively by the effector caspase, caspase 7. Cytosolic caspase 7 activity has been suggested to promote odontogenesis (Matalova et al. 2013; Svandova et al. 2018), yet nuclear caspase 7 disposition is considered to be an exclusive hallmark of apoptosis (Zhivotovsky et al. 1999; Kamada et al. 2005). For example, remodeling the chromatin micro-environment through targeted protein cleavage events is considered to be a conserved feature of apoptosis, as exemplified by effector caspase cleavage of scaffold attachment factor b1 (Lee et al., 2007), lamina-associated polypeptide 2α (Gotzmann et al., 2000), and SMARCA2 and SMARCA4 (Dudek et al., 2018). Nevertheless, our observations support a novel model whereby an effector caspase, caspase 7, targets a protein substrate (SATB2) to prime the nuclear matrix to engage differentiation, independent of cell death.

Presumably, during muscle cell differentiation the activation of caspase 7 is mediated by the same pathway that leads to the induction of caspase 3, via engagement of the mitochondrial intrinsic cell death pathway (Murry et al. 2008). The pattern of caspase 7 activation is remarkably similar to caspase 3 in differentiating myoblasts, which does support a common signaling origin. What is more speculative is whether a level of integration may exist between caspase 3 and caspase 7 activity, where targeting of their respective substrates is coordinated to drive the same biologic alteration. Given that SATB2 is a MAR protein, it is not unreasonable to suggest that caspase 7 cleavage of this protein may provide accessibility for CAD to target the genome and induce the requisite strand breaks during early stages of the differentiation program. Indeed, CAD does not target a specific DNA sequence, which suggests that the nuclease gains access to its genomic targets through an undefined structural change (Larsen and Sørenesen, 2017). However, our ChIP-seq analysis indicated that SATB2 was not enriched at a well-defined CAD target, the p21 promoter, and follow on ChIP PCR experiments validated this observation (Supplemental Figure 7). While this single observation does not preclude SATB2 shielding other undefined CAD targets, it does indicate that SATB2 and CAD have distinct genome reorganizing roles during muscle cell differentiation. As such, we favour the hypothesis that caspase 7 targeting of SATB2 is a temporally linked yet distinct biochemical step caspase mediated control of myogenesis. Finally, it is germaine to note that SATB2 is widely expressed and caspase signaling is an inductive cue for differentiation across multiple cell lineages and phyla. This convergence raises the possibility that caspase 7 cleavage of SATB2 may be a conserved mechanism to reprogram the nuclear architecture for cell differentiation.

## Acknowledgements

The authors would like to thank members of the Megeney laboratory and Michael Rudnicki for helpful discussion. We would also like to thank Lawrence Puente for the mass spectrometry analysis (Proteomics Core Facility, Ottawa Hospital Research Institute). This work was supported by grants from the Canadian Institutes of Health Research (L.A.M. and J.D.).

## Author Contributions

Conceptualization: L.A.M., J.D., G.D., and G.A. Formal analysis: R.A.V.B., M.H.A., and A.C. Investigation: R.A.V.B., M.H.A., and S.B. Writing – Original Draft: R.A.V.B. and M.H.A. Writing – Review and Editing: R.A.V.B., M.H.A., and L.A.M. Visualization: R.A.V.B. and M.H.A. Supervision: L.A.M. and J.D. Funding Acquisition: L.A.M.

## Supplemental Information

**Figure 1.**
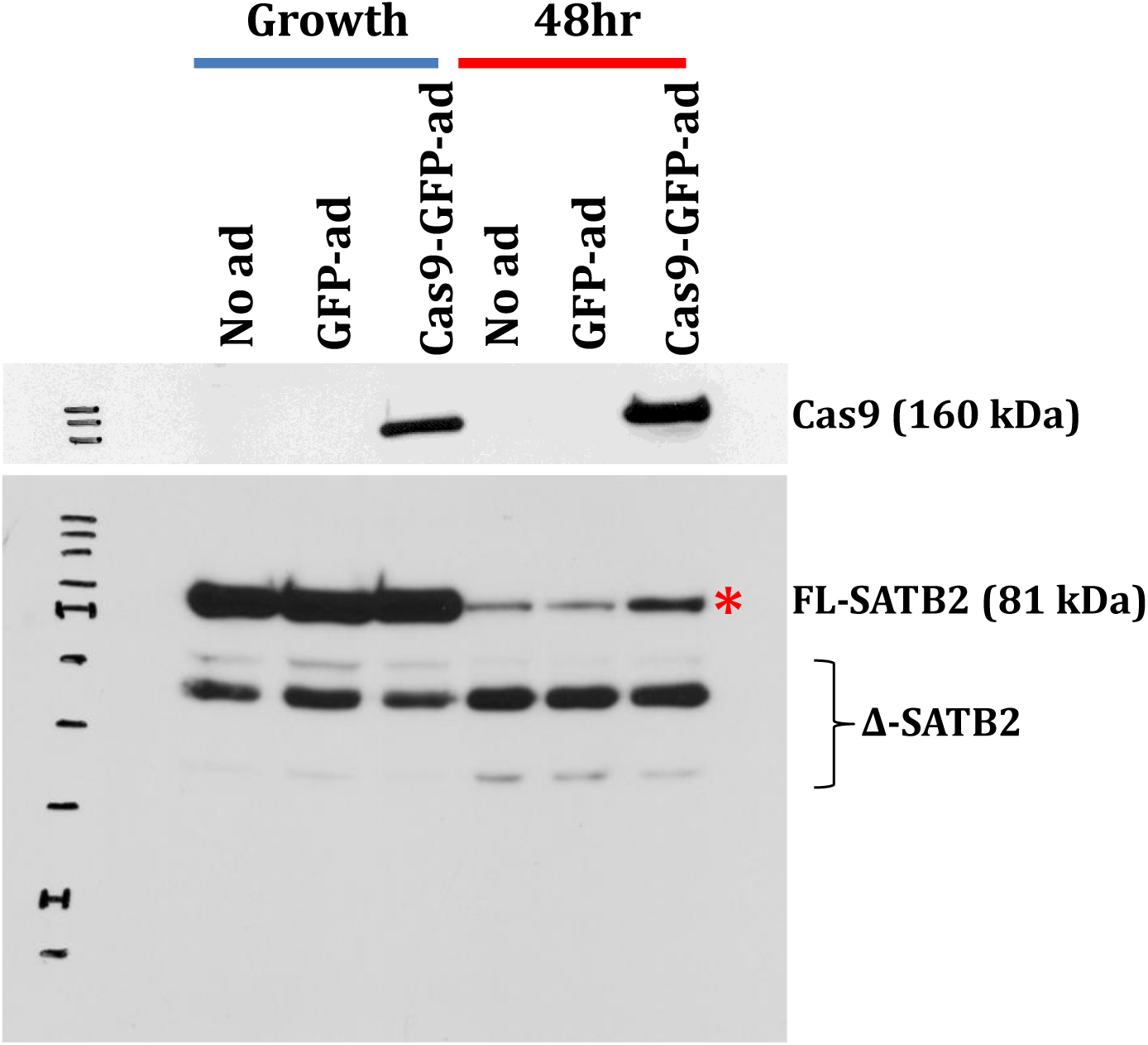
SATB2 expression following adenoviral Cas9 transfection into C2C12 myoblasts. SATB2 expression increased following Cas9-GFP-ad transfection as compared to no transfection and GFP-ad transfection controls (48 h differentiated C2C12 cells). * indicated the location of the full-length SATB2. Cas9 expression was confirmed in the Cas9-GFP-ad cells and is shown in a panel above the main blot.

**Figure 2.**
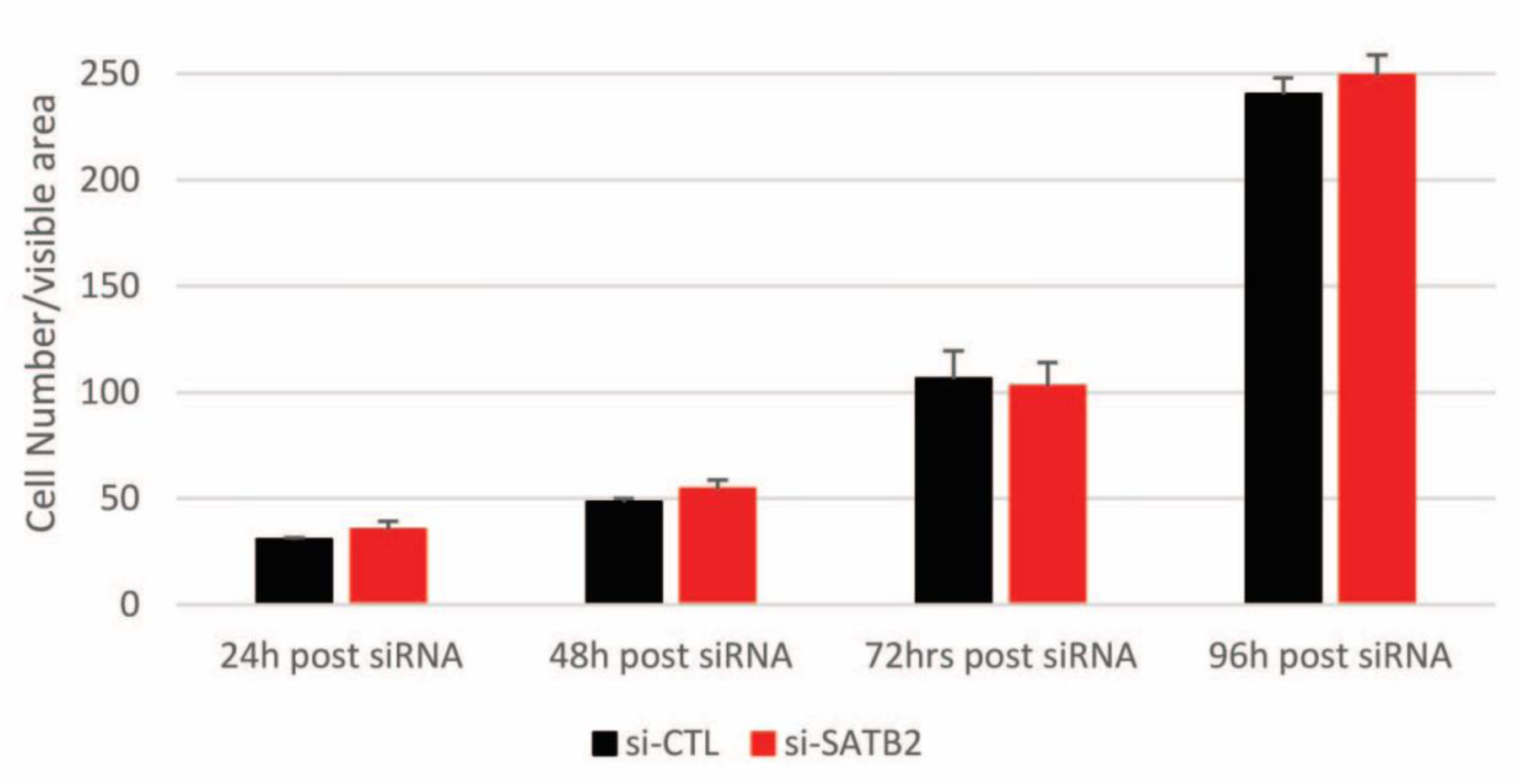
siRNA-mediated depletion of SATB2 does not affect C2C12 proliferation and survival. C2C12 cells were counted using bright field microscopic images under low magnification. Each individual cell in a field was marked and counted using ImageJ cell counter software. Data are means ± SEM for n = 3 independent determinations. No statistically significant difference in cell number was observed between siControl and siSATB2 treatments at each time point.

**Figure 3.**
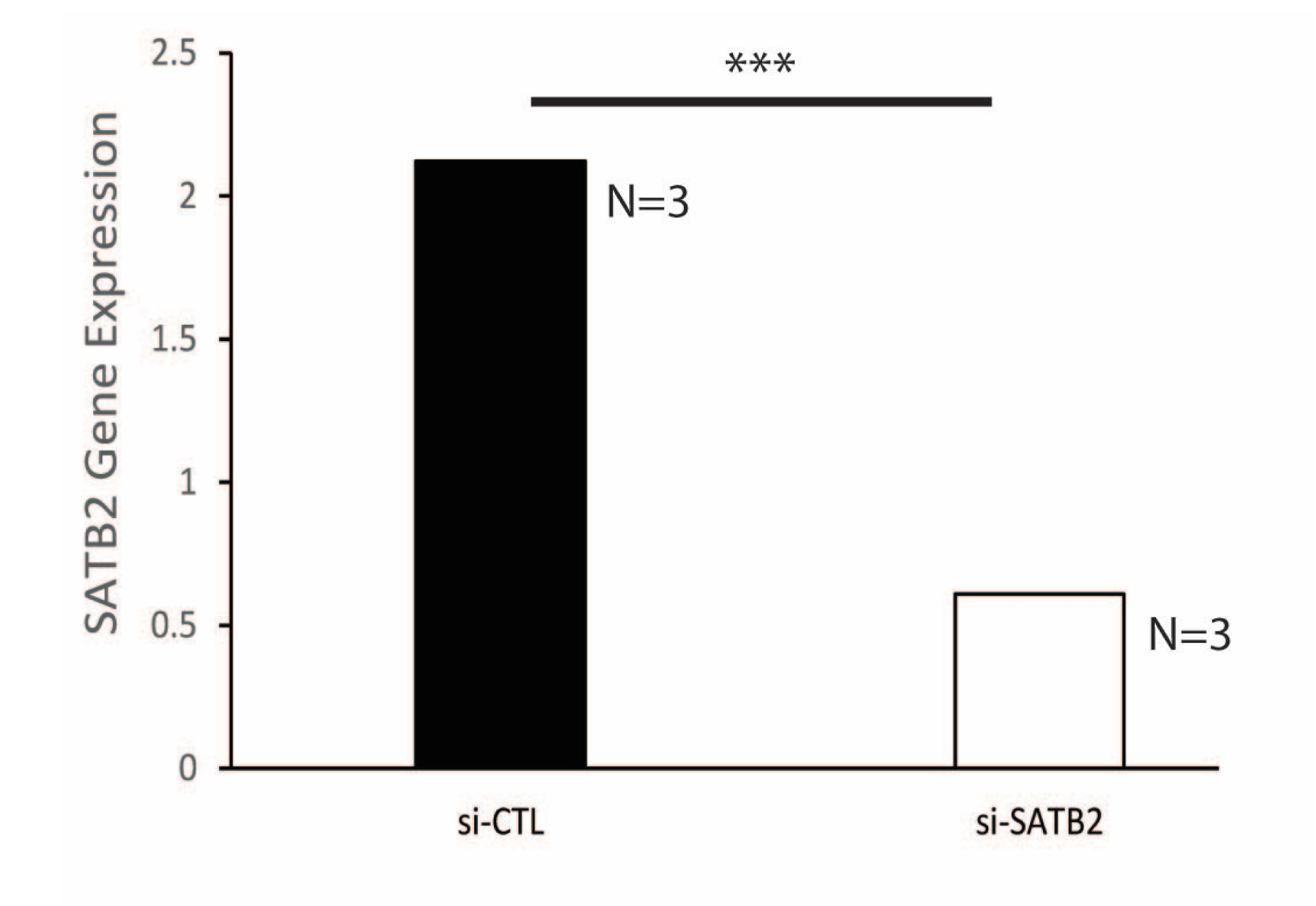
Validation of SATB2 downregulation following siSATB2 treatment via RNA-seq. The C2C12 cell samples were from the 24 h differentiated time point. *** indicates a significant difference between siControl and siSATB2 samples as determined by the Student’s *t*-test (p < 0.001).

**Figure 4.**
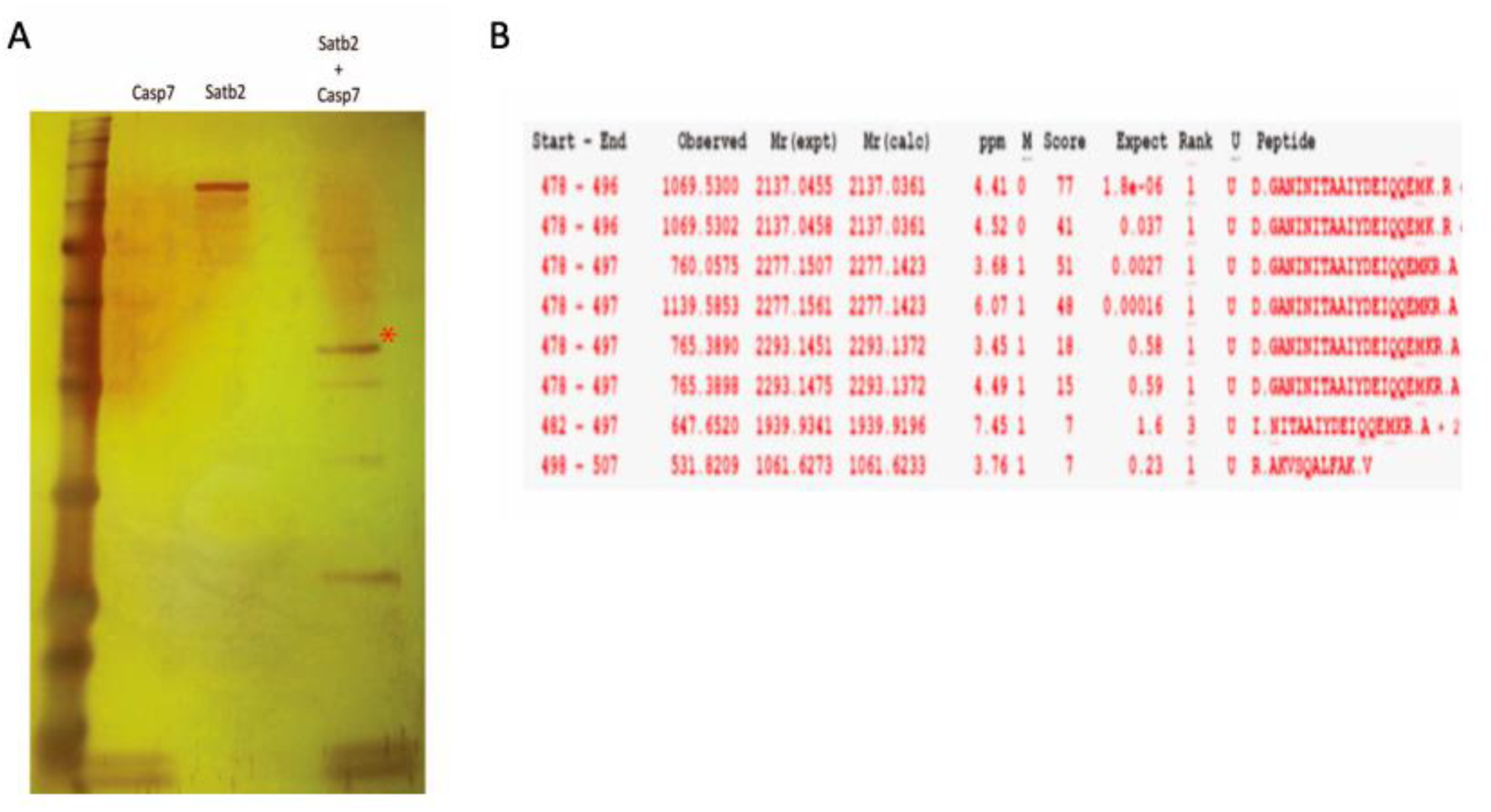
Identification of the caspase cleavage site within SATB2. (A) Silver stained gel of an in vitro caspase cleavage assay involving recombinant SATB2 and caspase 7. * indicates the major cleavage fragment that was analyzed by mass spectrometry and included the D477 cleavage site. (B) Mass spectrometry analysis of the major cleavage fragment SATB2, identified with an * in (A).

**Figure 5.**
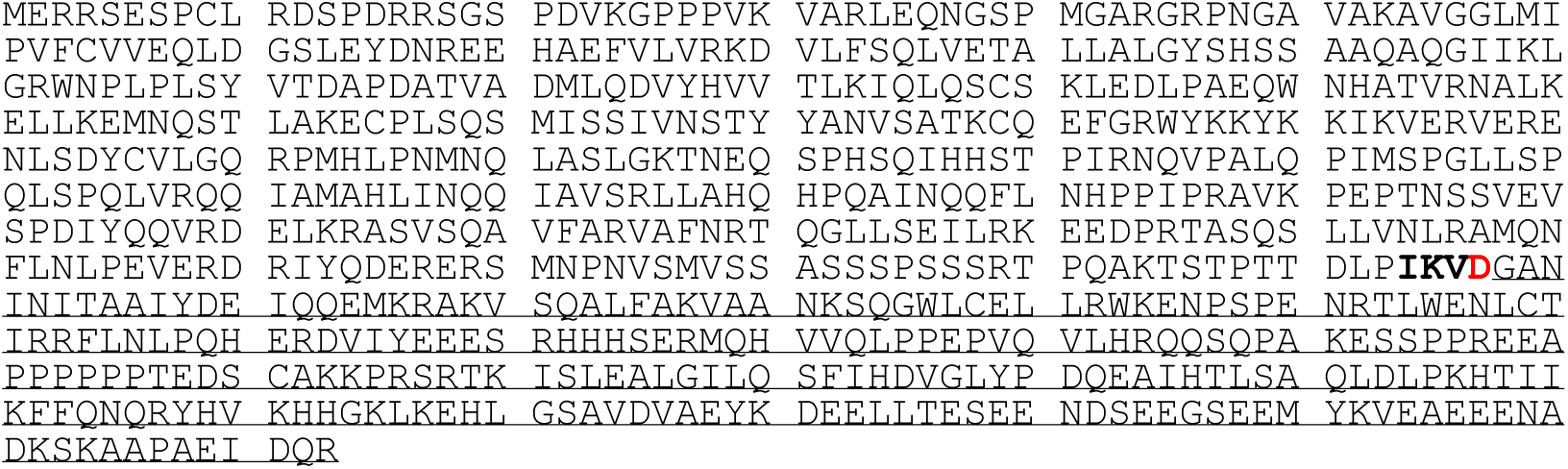
The amino acid sequence of *Mus musculus* SATB2, with the major cleavage fragment underlined and the caspase cleavage fragment bolded. The red ‘D’ indicates the aspartic acid residue where cleavage occurs.

**Figure 6.**
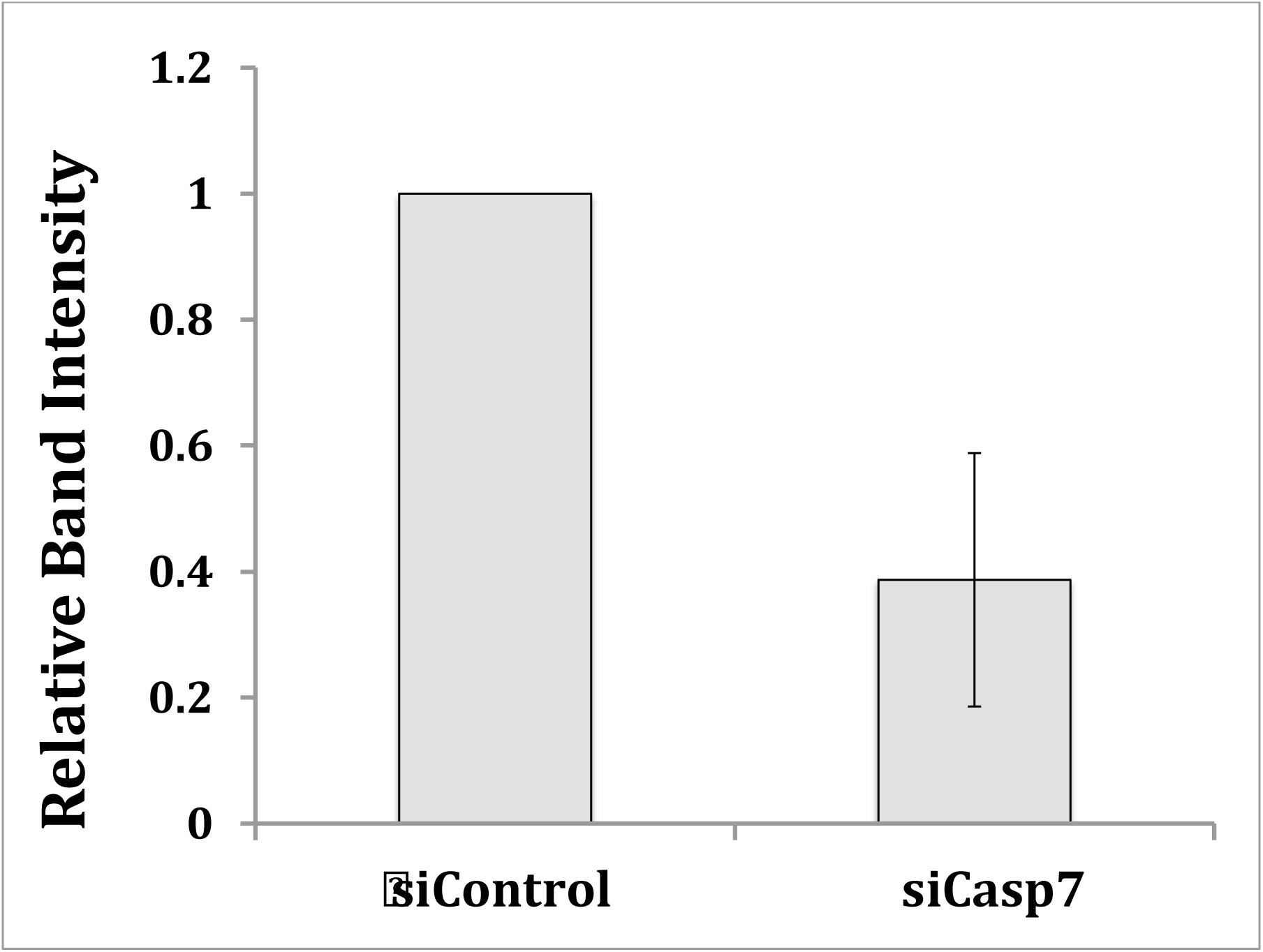
Relative caspase 7 expression following knockdown with siCasp7 in differentiating C2C12 myoblasts. Data are means ± SEM, n = 3 independent determinations. * indicates a statistically significant difference versus the siControl condition using the Student’s *t*-test, p < 0.05.

**Figure 7.**
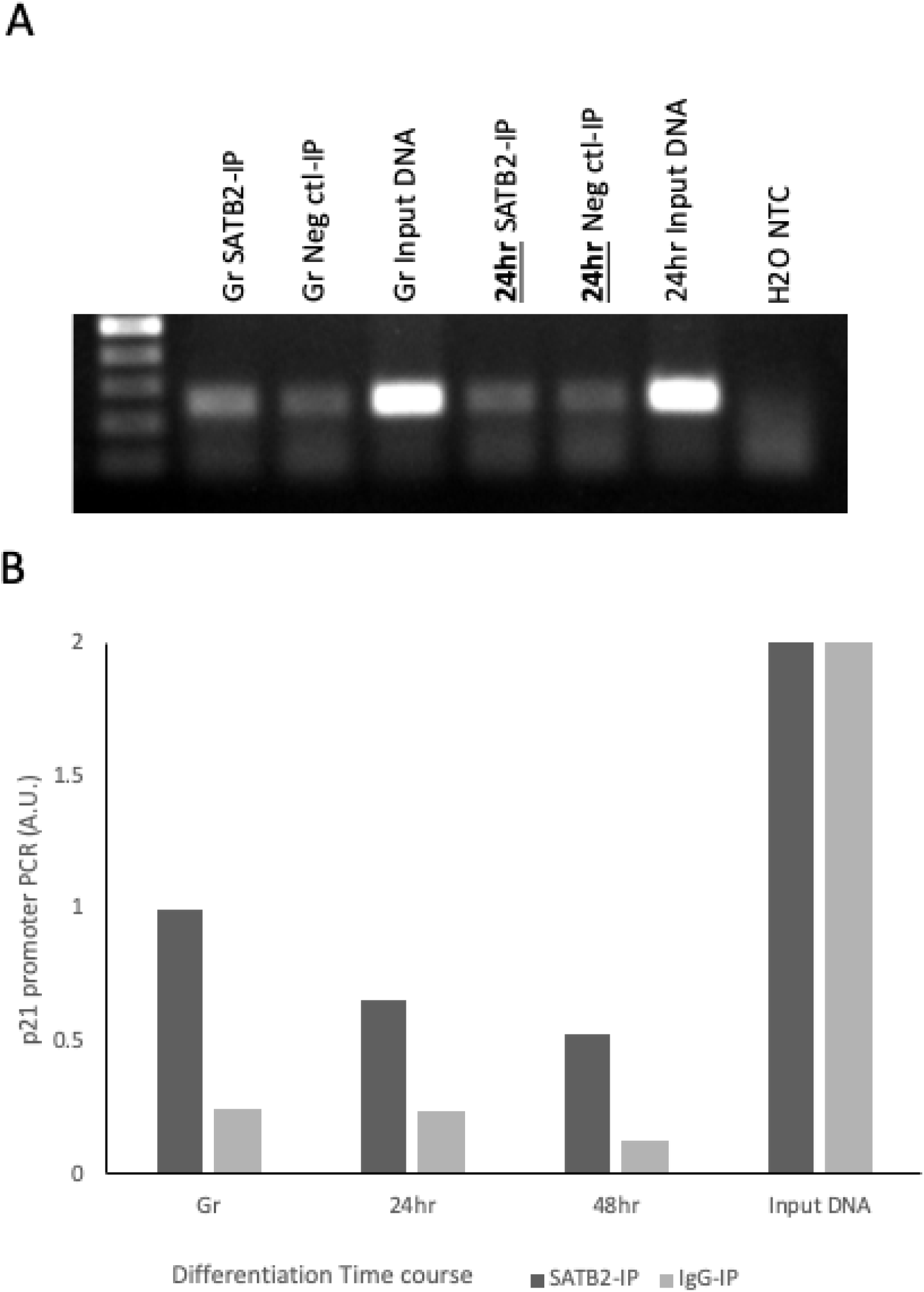
Enrichment of SATB2 at the *p21* locus during C2C12 differentiation. Chromatin immunoprecipitation (ChIP) using an anti-SATB2 antibody indicted that SATB2 was not significantly enriched at the *p21* locus during cell proliferation or differentiation. (A) Representative PCR amplification of *p21* following ChIP using a SATB2 antibody. The negative control was ChIP using a mouse IgG. (B) Quantification of ChIP PCR results for the *p21* locus.

## Supplementary Tables

**Table 1.**
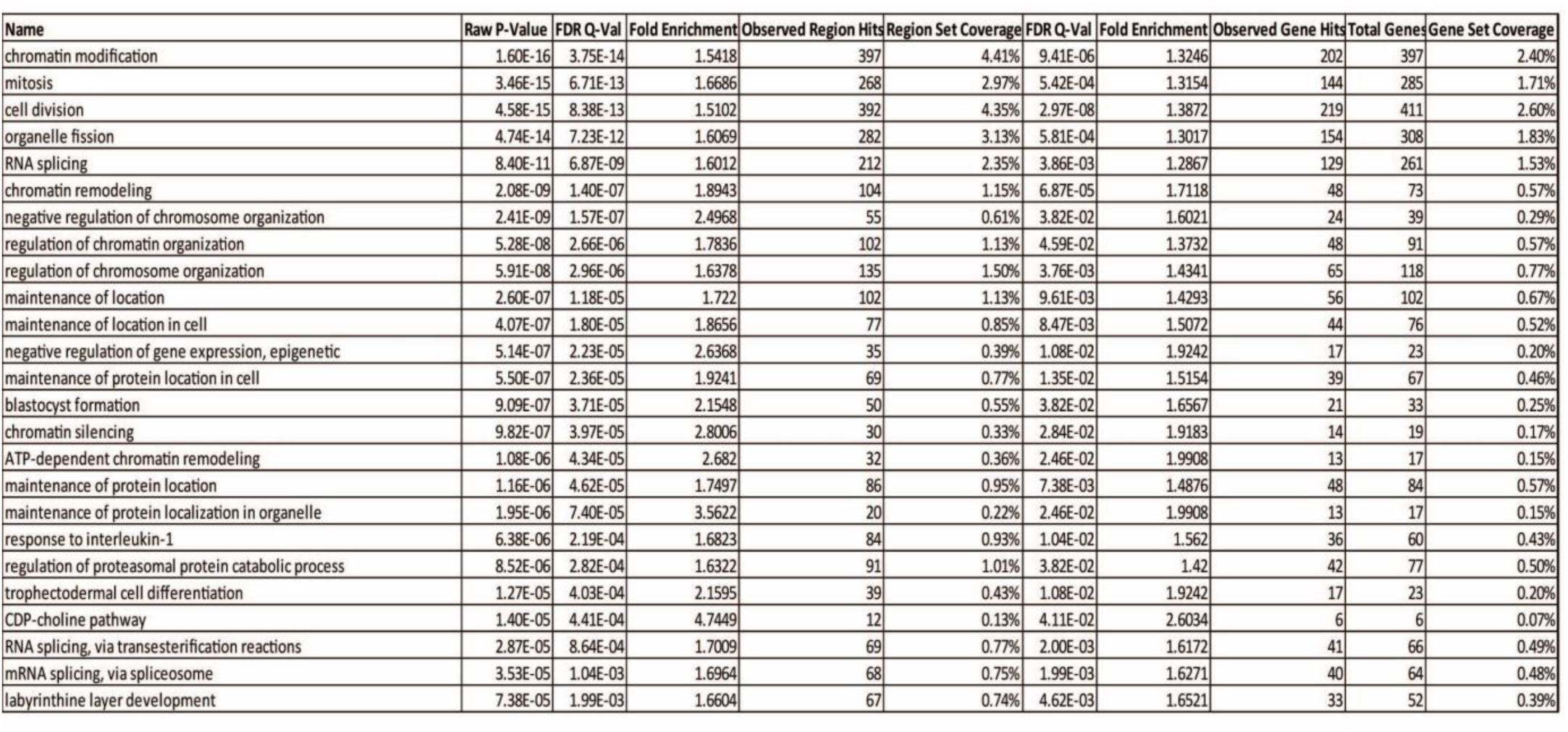
Top 25 Gene Ontology biological processes identified within the SATB2-ChIP-seq dataset.

**Table 2.**
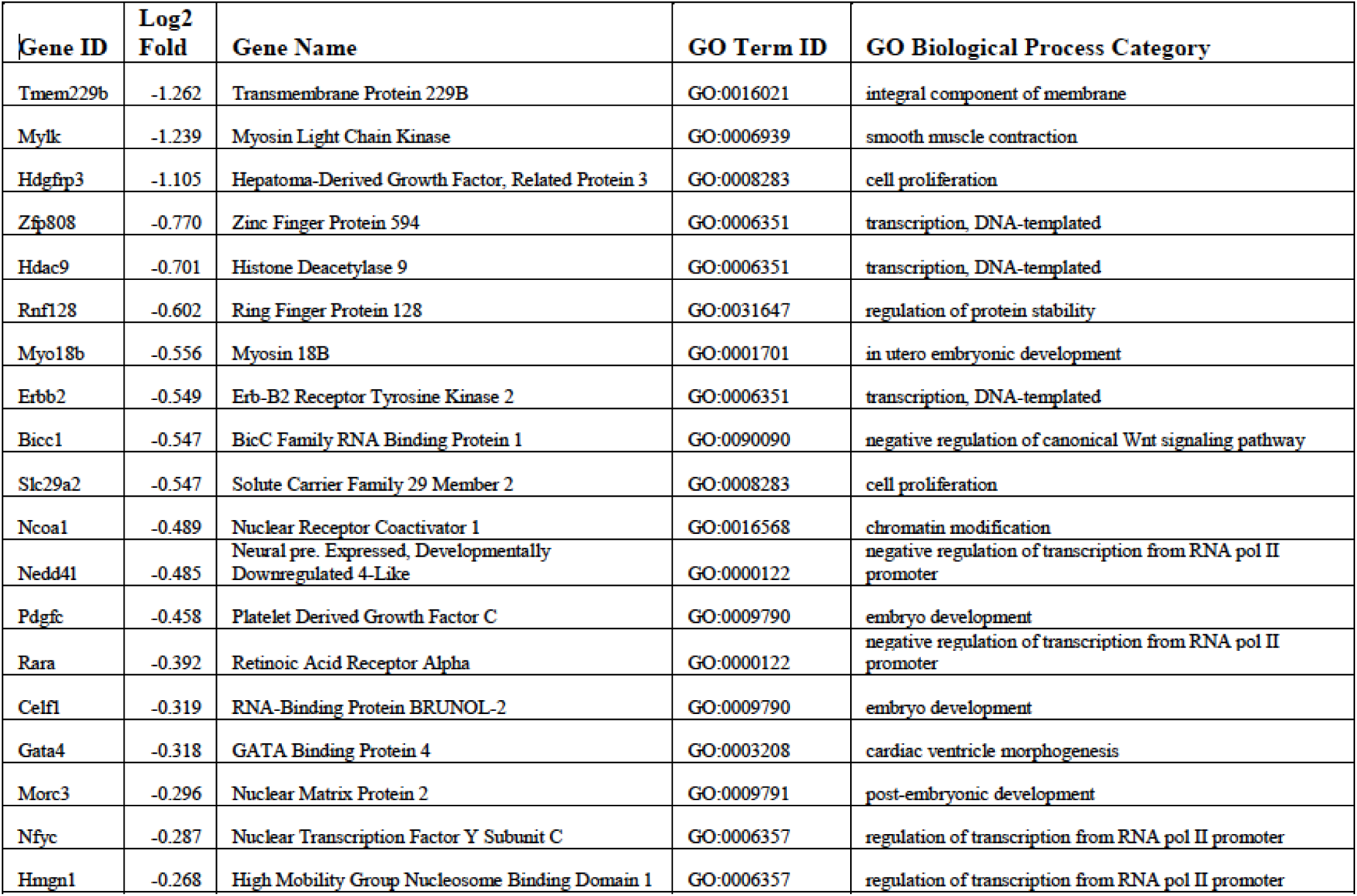
RNA-seq-identified genes that were downregulated following *Satb2* depletion in C2C12 cells. The genes shown here were also part of the ChIP-seq dataset.

**Table 3.** RNA-seq-identified genes that were upregulated following *Satb2* depletion in C2C12 cells. The genes shown here were also part of the ChIP-seq dataset.

| Gene ID | Log2 Fold | Gene Name | GO Term ID | GO Biological Process Category |
| --- | --- | --- | --- | --- |
| Taf9 | 0.274 | TATA-Box Binding Protein Associated Factor 9 | GO:0016568 | chromatin modification |
| Smarca4 | 0.279 | SWI/SNF rel. Matrix Assoc. Actin-dep. Regulator Chromatin A4 | GO:0016568 | chromatin modification |
| Hivep1 | 0.295 | Positive Regulatory Domain II Binding Factor 1 | GO:0006366 | transcription from RNA pol II promoter |
| Qk | 0.300 | Quaking Homolog, KH Domain RNA Binding | GO:0008380 | RNA splicing |
| Gtf3c3 | 0.302 | General Transcription Factor IIIC Subunit 3 | GO:0006351 | transcription, DNA-templated |
| Wnt9a | 0.309 | Wnt Family Member 9A | GO:0008285 | negative regulation of cell proliferation |
| Med21 | 0.339 | Mediator Of RNA Polymerase II Transcription Subunit 21 | GO:0006357 | regulation of transcription from RNA pol II promoter |
| Cdk8 | 0.357 | Cyclin Dependent Kinase 8 | GO:0051726 | regulation of cell cycle |
| Rad54l2 | 0.413 | Helicase ARIP4 | GO:0005634 | nucleus |
| Sorbs2 | 0.449 | Arg Binding Protein 2 | GO:0007015 | actin filament organization |
| Runx2 | 0.487 | Runt Related Transcription Factor 2 | GO:0001501 | skeletal system development |
| Kat2b | 0.489 | Histone Acetyltransferase KAT2B | GO:0016568 | chromatin modification |
| Daam1 | 0.515 | Dishevelled Associated Activator Morphogenesis 1 | GO:0030036 | actin cytoskeleton organization |
| Ddx19b | 0.533 | DEAD-Box Helicase 19B | GO:0006406 | mRNA export from nucleus |
| Ckap2 | 0.597 | Cytoskeleton Associated Protein 2 | GO:0045944 | positive regulation of transcription from RNA pol II promoter |
| Bdnf | 0.633 | Brain Derived Neurotrophic Factor | GO:0031547 | brain-derived neurotrophic factor receptor signaling pathway |
| Zfp521 | 0.711 | Zinc Finger Protein 521 | GO:0006351 | transcription, DNA-templated |
| Cdk6 | 0.852 | Cyclin Dependent Kinase 6 | GO:0000082 | G1/S transition of mitotic cell cycle |
| Rcan2 | 0.870 | Regulator Of Calcineurin 2 | GO:0070884 | regulation of calcineurin-NFAT signaling cascade |

## Materials and Methods

### Mice and in vivo procedures

All mice were housed and treated at the University of Ottawa Animal Care and Veterinary Services. Mice used in our studies were housed and cared for according to Canadian Council on Animal Care (CCAC) guidelines and University of Ottawa Animal Care Committee protocols.

In order to determine the effects of SATB2 removal from muscle satellite cells, *Pax7/CreER Satb2_fl/fl_* and Satb2_fl/fl_ (control) mice were fed a diet supplemented with tamoxifen (40 mg/kg body weight; Envigo) at 3 weeks old. These mice were monitored weekly for evidence of potential effects of SATB2 ablation from muscle stem cells. After 8–10 months, three *Pax7/CreER Satb2_fl/fl_* and *Satb2_fl/fl_* mice (female) were euthanized by CO_2_ asphyxiation and cervical dislocation. These mice were then assessed for any structural abnormalities, and their tibialis anterior muscle was excised and fixed in 10% neutral buffered formalin. After 24 h, the samples were then placed in 70% ethanol and then paraffin embedded and cut into 4 μm sections. To determine muscle fiber areas, sections were stained with hematoxylin and eosin (H&E) and assessed under an Observer A1 microscope (Zeiss). Muscle fiber area was determined using the ImageJ software. To investigate the percentage of Pax7-expressing satellite cells, sections were stained with a Pax7 antibody, as well as just the secondary antibody as a control.

### Cell culture

Cells of the immortalized mouse cell line, C2C12, were grown on non-collagen coated cell culture plates in Dulbecco’s Modified Eagle’s medium (DMEM) with 10% fetal bovine serum (FBS) and 1% penicillin/streptomycin (growth medium). When confluent, cells were differentiated in DMEM supplemented with 2% horse serum and 1% penicillin/streptomycin (differentiation medium).

### Protein extraction and western blotting

C2C12 cells were lysed using a modified RIPA buffer (50 mM Tris-HCl, pH 7.4, 150 mM NaCl, 1 mM ethylenediaminetetraacetic acid (EDTA), 1% NP-40, 1% glycerol, and a cocktail of protease inhibitors) for whole cell extracts, or a modified NE-PER Nuclear and Cytoplasmic Extraction Kit, as per the manufacturer’s instructions (Thermo Scientific). Extracted proteins were then separated via SDS-PAGE and then transferred to a 0.45 μM polyvinylidene fluoride (PVDF) membrane (Millipore) on a TRANS-BLOT SD apparatus (Bio-Rad). Membranes were blocked with Tris-buffered saline plus 0.1% Tween-20 (TBST) containing 5% skim milk for 1 h at room temperature. Membranes were then incubated overnight at 4 °C with primary antibody made in blocking solution. The primary antibodies used in this study included mouse SATB2 (ab51502; Abcam; 1:1000), rabbit caspase 7 (9491S; Cell Signaling; 1:1000), mouse myosin heavy chain (DSHB; MHC; 1:250), and mouse glyceraldehyde 3-phosphate dehydrogenase (#2118; GAPDH; 1:4000).

### Immunofluorescence

C2C12 cells were cultured and fixed in paraformaldehyde on 25 mm coverslips at the desired time points. The cells were then incubated for 10 min at room temperature with a permeabilization solution containing 0.5% Triton-X 100 in PBS. Subsequently, the cells were incubated for 1 h with a blocking solution consisting of 5% horse serum in PBS. After blocking, the cells were incubated for 2 h at room temperature or overnight at 4 °C in primary antibody that was reconstituted in blocking solution. The primary antibodies used were mouse SATB2 (1:50; ab51502, Abcam), rabbit HP1α rabbit (1:200; #2616, Cell Signaling), rabbit Gapdh (1:400; #2118, Cell Signaling), rabbit desmin (1:400; ab15200, Abcam), and mouse MHC (1:50; DSHB). After primary antibody incubation, the cells were washed three times in PBS and incubated in secondary antibody (2 mg/mL Alexa Fluor 488, Invitrogen, 1:1500, 2 mg/mL Alexa Fluor 594, Invitrogen, 1:1500; 2 mg/mL Alexa Fluor 568, Invitrogen, 1:1000) diluted in PBS for 1.5 h at room temperature. After incubation, the cells were washed 2× in PBS and then counterstained with 4′,6-diamidino-2-phenylindole dihydrochloride (DAPI; 1:10 000; Sigma) for 10 min at room temperature. After incubation, the cells were washed 2× in PBS and the coverslips were mounted on microscope slides using Dako Fluorescent Mounting Medium. Cells were then visualized using a Zeiss Observer Z1 inverted fluorescence microscope. All images were developed using AxioVision 4.8 software. The quantification of nuclear SATB2 was performed by densitometry analysis using ImageJ/Photoshop C3 software.

### Chromatin immunoprecipitation (ChIP) assay

Cells were grown on 15 cm plates and allowed to reach 100% confluence before either being collected or switching to differentiation media and allowed to incubate until the desired time point was achieved. Twenty million cells per time point were used for each ChIP sample. Cells were fixed for 10 min using 1% formaldehyde in DMEM. Cell fixation was quenched by removing the fixation solution, rinsing the plates with PBS, and then pouring a solution of 0.125 M glycine in PBS onto the cells and incubating for 5 min. Subsequently, plates were washed 2× with PBS and then cells were scraped from the plates, pelleted, and stored at −80 °C until used.

Given that SATB2 is a nuclear matrix attachment protein, it required an alternative protocol for chromatin shearing than standard preparations. First, a commercial hypotonic solution (Active Motif) was used to lyse cellular membranes but retain intact nuclear membranes. We subsequently performed additional lysis using a hand held glass Dounce homogenizer. After centrifugation, the supernatant was discarded and the pelleted nuclei were treated with Active Motif’s Pro-enzymatic Digestion Nuclear Extraction Solution. After incubation at 37 °C for 5 min, we supplemented the reaction with 1 U micrococcal nuclease (New England Biolabs) and allowed it to incubate for 10 min. We then supplemented the reaction with SDS to a final volume 2% SDS-chromatin solution. We proceeded to the second stage of shearing using sonication via a Covaris M220 Focused-Ultrasonicator instrument, setting the parameters to produce sheared DNA of 200 bp. After sonication, we performed a final centrifugation at 16 000 × *g* at 4 °C for 20 min and collected the supernatant containing the sheared chromatin.

We performed immunoprecipitations with 5 μg 1°Ab (Target: SATB2, Abcam, positive control: RNA pol II, Active Motif, negative control: mouse IgG, Santa Cruz) using magnetic beads (Active Motif) diluted in commercial ChIP-buffer (Active Motif) overnight at 4 °C. Captured chromatin was isolated using magnetic stands to pulldown the beads. Bound DNA was subsequently eluted with commercial elution, reverse cross-linking, protein digestion, and RNA digestion solutions (Active Motif). The resulting samples of DNA were further purified using phenol:chloroform extraction procedures. Final DNA was quantified using a NanoDrop spectrophotometer, with a 25 μL final volume having concentrations of 75–100 ng/μL.

### ChIP-sequencing and bioinformatics

For genome-wide analysis, immunoprecipitated DNA was amplified and 75 bp single read sequencing was performed on an Illumina HiSeq 2500 at the Next-Generation Sequencing Facility at The Centre for Applied Genomics in The Hospital for Sick Children (Toronto, Ontario, Canada). The data were summarized and basic comparisons performed using the Excel spreadsheet program (Microsoft). Reads were aligned to the NCBI build 38 (UCSC mm10, Dec/2011) genome from the UCSC genome browser using the default options. Data were summarized and basic comparisons performed using MACS version 2.1.0.20140616, and Gene Ontology (GO) term annotation was performed using GREAT v3.0.0 available online from the Bejerano lab at Stanford University (McLean et al, 2010).

### RNA-sequencing and bioinformatics

Total RNA was isolated using an RNeasy Kit (QIAGEN) using an on-column DNase digestion (RNase-Free DNase Set, QIAGEN) to avoid genomic DNA contamination. Library preparation and 126-bp paired-end RNA-seq was performed by the Next-Generation Sequencing Facility at The Centre for Applied Genomics in The Hospital for Sick Children. RNA integrity was assessed using the Bioanalyzer platform (Agilent Technologies, Inc.). Sequencing was performed using standard procedures for the Illumina HiSeq 2500 platform. Gene expression quantification was performed using CuffDiff 2 software (Trapnell et al 2013). Data were summarized and basic comparisons were performed using the Excel spreadsheet program (Microsoft).

### Caspase cleavage assays

Recombinant SATB2 protein (250–500 ng; Abnova) and recombinant active caspase 3 (0.5 μg; Chemicon) or recombinant active caspase 7 (0.5 μg; Biovision) were incubated for 3 h in cleavage assay buffer (50 mM Hepes, pH 7.5, 0.1 M NaCl, 10% (v/v) glycerol, 0.1% Chaps, 10 mM dithiothreitol) containing either dimethyl sulphoxide (DMSO) or z.DEVD.fmk (20 μM; BioVision) as indicated. Reactions were stopped by the addition of Laemmli sample buffer, and subjected to sodium dodecyl sulphide polyacrylamide gel electrophoresis (SDS/PAGE). Mass spectrometry was performed at the Ottawa Hospital Research Institute Proteomics Core Facility (Ottawa, Canada). MASCOT 2.3.01 software (Matrix Science) was used to infer peptide and protein identities from the mass spectra.

### Caspase inhibition assays

For caspase 3/7 inhibition, cultured C2C12s were pre-treated with either 15 μM z-DEVD-fmk (DEVD) from BioVision or 15 μM DMSO from Sigma for 2 h at 37 °C. After pre-treatment, the cells were induced to differentiate using low serum media or continued in growth media both containing 15 μM DEVD or DMSO as a vehicle-only control. The inhibition or control media was changed every 48 h until the end of the time course. Cells were collected at the predetermined time points and analyzed as described.

### siRNA-mediated depletion of SATB2 and caspase 7 gene expression

siRNA duplexes were used to suppress SATB2 and caspase 7 gene expression in C2C12 cells. C2C12 cells were transfected at 25% confluence with 10 nM siRNA (siSATB2, siCasp7, or siControl) and the Lipofectamine RNAiMAX reagent, as directed by the manufacturer’s protocol (Invitrogen). After an overnight incubation, fresh media was added onto the cells and the cells were re-transfected. This continued until the cells reached 100% confluence, after which differentiation media was added to the cells. Media change and re-transfection occurred every 24 h throughout the indicated time course. Cells were collected or used in subsequent experiments at a growth time point and 24, 48, 72, and 96 h of differentiation.

### Statistical Analysis

Statistical analysis of three or more data sets was performed using one-way analysis of variance (ANOVA). For comparison between two sample sets, an unpaired, 2-tailed Student’s *t*-test was performed. p < 0.05 was considered statistically significant.

